# Explaining tip-of-the-tongue experiences in older adults: The role of brain-based and cardiorespiratory fitness factors

**DOI:** 10.1101/2023.12.08.570799

**Authors:** Foyzul Rahman, Kamen A. Tsvetanov, Jack Feron, Karen Mullinger, Kelsey Joyce, Ahmed Gilani, Eunice G. Fernandes, Allison Wetterlin, Linda Wheeldon, Samuel J.E. Lucas, Katrien Segaert

## Abstract

Cognitive decline associated with healthy ageing is multifactorial: brain-based and lifestyle factors uniquely and jointly contribute to distinct neurocognitive trajectories of ageing. To evaluate existing models of neurocognitive ageing such as compensation, maintenance, or reserve, we explore how various known brain-based and cardiorespiratory fitness factors intersect to better understand cognitive decline. In a pre-registered study (https://osf.io/6fqg7), we tested 73 healthy older adults aged 60—81 (*M* = 65.51, *SD* = 4.94) and collected neuroimaging (functional, structural, and perfusion MRI), cardiorespiratory fitness, and cognitive data to investigate a prominent challenge for older adults: word-finding failures. fMRI signal was recorded while participants responded to a definition-based tip-of-the-tongue task, T1-weighted imaging estimated grey matter volume, and cerebral blood flow was indexed using multi-delay pseudo-continuous arterial spin labelling. Commonality analyses were used to analyse these multi-domain data (neuroimaging, cardiorespiratory fitness, language skills, demographic characteristics) and uncover associations between predictors in explaining age-related tip-of-the-tongue rates. Commonality analyses revealed that functional activation of language networks associated with tip-of-the-tongue states is in part linked with age and, interestingly, cardiorespiratory fitness: the combination of higher cardiorespiratory fitness and functional recruitment in some older adults offsets part of the age-related variance in tip-of-the-tongues. Moreover, age-associated atrophy and perfusion in regions other than those showing functional differences accounted for variance in tip-of-the-tongues. Our findings can be interpreted in the context of the classic models of neurocognitive ageing, suggesting compensation. Brain health indices in concordance with cardiorespiratory fitness can provide a more holistic explanation of individual differences in age-related cognitive decline.

**Highlights:**

- Word-finding problems are linked to brain health and cardiorespiratory fitness (CRF)
- Brain activity linked to word-finding failures is modulated by CRF and age
- Distinct contribution of structure and perfusion also associated with word-finding
- Linking brain and CRF factors provides better account of age-related cognitive decline

## Introduction

The proportion of older adults in the global population is rapidly increasing, a trend most pronounced in high-income countries (WHO, 2022). Despite this, there are considerable gaps in our understanding of how we can promote maintenance of cognitive function in later life. Even in the absence of pathology in older age, cognitive decline—including difficulties with memory, processing, and communication skills—can limit everyday functioning and restrict independent living (Clouston et al., 2013; Njegovan et al., 2001). One prominent example of this is the increasing difficulty in older age with word finding and a related phenomenon known as tip-of-the-tongue states. These are impermanent failures to retrieve and produce intended words; lapses where we cannot call upon a word that we know. Virtually all speakers experience tip-of-the-tongue states during regular, everyday conversation but they are of particular significance to older adults, who regard them as their most frustrating and troubling cognitive failure (Ossher, Flegal, & Lustig, 2013). Existing neurocognitive accounts show that tip-of-the-tongue states are associated with structural and functional changes in the brain (Huijbers et al., 2017; Resnik et al., 2014; Shafto et al., 2007; 2010), and behavioural performance studies have found that cardiorespiratory fitness levels modulate individuals’ abilities to resolve tip-of-the-tongue states (Segaert et al., 2018). Indeed, more generally, cardiorespiratory fitness has been shown to be an important factor in successful neurocognitive ageing across studies of brain function (Kawagoe, Onoda, & Yamaguchi, 2017; Prakash et al., 2011), brain structure (Colcombe et al., 2003; Szabo et al., 2011), and cognitive performance (Barnes et al., 2003; Hayes, Forman, & Verfaellie, 2016). However, returning to the issue of tip-of-the-tongue states, an integrated explanation of these highly prevalent cognitive failures, which includes cardiorespiratory fitness as a variable in addition to brain-based measures, is currently lacking. To address this gap, it is essential to examine how fitness and brain-based factors, including function, structure, and perfusion, contribute to age-related differences in tip-of-the-tongue states. These insights need to be aligned with established models of cognitive ageing, which we turn to next.

Key models of healthy ageing centre around mechanisms of maintenance, compensation, or reserve and typically assume contributions of both genetic and lifestyle influences, such as an individual’s fitness level, to the neurocognitive mechanisms of healthy ageing (Cabeza et al., 2018). Models of maintenance are established on maintaining or returning neural resources to their former levels in response to the typical ‘wear and tear’ associated with non-pathological ageing (Habeck et al., 2017; Nyberg et al., 2012). As such, fitness levels are predicted to promote brain maintenance by preserving youthful brain structure and cerebrovasculature, potentially supporting brain function and cognition through grey matter preservation and optimal cerebral perfusion. Models of compensation entail functional recruitment of additional resources to meet high cognitive demands when there is an insufficiency in neural resources (e.g., brain atrophy or hypoperfusion) (Reuter-Lorenz & Cappell, 2008). Typically, observations of increased recruitment of particular neural regions in older (vs younger) adults are taken as evidence for compensation, with the presumption that these increases offset neural decline elsewhere in the brain (Knights et al., 2024). Nevertheless, a key assumption with this model is that the degree of compensatory involvement is directly related to the degree of cognitive performance, an assumption which is not tested directly in most previous studies. Lastly, models of reserve typically posit the idea that fitness, education, or crystallised intelligence (which are accepted to be proxies for reserve) can contribute to accumulated neural reserve, which can be structural and/or functional and develops over time (e.g., through childhood or early adulthood) (Stern et al., 2020; Tucker & Stern, 2011; Whalley et al., 2004). This reserve is thought to protect against age-related neural pathology. Consequently, proxies of cognitive resilience are expected to correlate with brain functional resources involved in specific cognitive processes, rather than structural differences.

As alluded to above, tip-of-the-tongue states signify difficulties with word retrieval. When one produces a spoken word, a lexical representation is selected based on its match to semantic information, and the associated phonology is accessed (e.g. Abrams & Davis, 2016; Levelt, 2001). For most people, most of the time, this process is effortless. However, in older age, connections throughout the lexical system are weakened, and the transmission from meaning to phonology is thought to be compromised, resulting in increased occurrences of tip-of-the-tongue states (Burke, MacKay, Worthley, & Wade, 1991). Different loci for tip-of-the-tongue states have been suggested. One proposal is that they occur due to failed lexical selection at the level intermediate between semantic and phonological representations (Gollan & Acenas, 2004); another is that they are due to failed word-form selection following successful lexical retrieval (e.g., Burke et al., 1991). Importantly for our study, the processes contributing to this type of cognitive failure thus involve contributions that are distinct from general cognitive slowing (Salthouse & Mandell, 2013). Complementing these psycholinguistic accounts, several neuroimaging studies have demonstrated neural markers associated with tip-of-the-tongue states: lower grey matter density in brain areas implicated in phonological production, in particular the left insula is related to greater frequencies of tip-of-the-tongue states (Shafto et al., 2007), and reduced functional activation in this same brain region has been demonstrated for older adults (Shafto et al., 2010). Several studies have demonstrated distinct neural signatures for successfully recollected responses compared to tip-of-the-tongue states (Galdo-Alvarez, Lindın, & Diaz, 2009), including recruitment of the wider lateral prefrontal cortex (Huijbers et al. 2017). The increased prevalence of tip-of-the-tongue states in older adults may be linked to age-related decline in key brain regions involved in word retrieval. Structural and functional changes in the posterior middle temporal gyrus and anterior temporal lobes can impair access to lexical representations and semantics, respectively (Binder et al., 2003; Visser, Jefferies, & Lambon Ralph, 2010). Changes in area Spt may disrupt phonological processing (Hickok et al., 2003) and changes in the inferior frontal cortex may impact domain-general processes (Fedorenko, 2014), such as cognitive control during retrieval (Meinzer et al., 2012). Declines in brain perfusion and connectivity, potentially mitigated by higher cardiorespiratory fitness, may further contribute to retrieval difficulties (Callow & Smith, 2023; Galiano et al., 2020). Modifiable lifestyle factors, such as fitness, have been shown to decrease the probability of experiencing tip-of-the-tongue states in healthy older adults (Segaert et al., 2018). However, these individual and isolated accounts have led to a fragmented picture; how different measures of brain health (structural, functional, and vascular) intersect with each other as well as with cardiorespiratory fitness in determining age-associated cognitive decline remains largely an underexplored area.

The present research aims for a more comprehensive account of how ageing leads to cognitive lapses, such as tip-of-the-tongue states. A multi-modal approach, integrating brain-based and lifestyle assessments, is necessary to examine how each factor uniquely and collectively explains this phenomenon. A suitable approach to address this is commonality analysis (Nimon & Oswald, 2013; Nimon et al., 2008), as it partitions explained variance into unique and shared contributions. Partitioning variance subsequently provides a clearer understanding of predictor relationships compared to traditional regression analysis, which can be distorted by multicollinearity, especially when age is a factor (Nimon et al., 2008; see Wu et al. 2023 for a recent and comprehensive example of applying commonality analysis to neuroimaging data).

Using commonality analysis, we assessed the unique and shared effects of brain function (networks revealed by task-based fMRI), grey matter volume, cerebral perfusion, cardiorespiratory fitness via a gold-standard lab-based incremental treadmill test, education and age to explain tip-of-the-tongue behaviour. Together, this approach allowed us to test the following predictions, associated with key neurocognitive ageing models: (1) Brain maintenance may be indicated by shared signals among grey matter, perfusion, and brain activity associated with tip-of-the-tongue incidence. This supports the notion that preserving youthful brain structure and function is key to maintaining cognitive health. (2) Compensation may be reflected by unique associations between activity in newly recruited regions and performance during tip-of-the-tongue states, not shared with grey matter (i.e., this functional recruitment would offset atrophy elsewhere). (3) Evidence for reserve could be reflected in shared coefficients between education or crystallised intelligence and brain activation with tip-of-the-tongue performance. (4) Importantly, if fitness levels also share signals with indicators of either maintenance, compensation or reserve, it suggests that fitness promotes these adaptive brain conditions.

As such, different neurocognitive models of ageing, centring around maintenance, compensation, or reserve, make different predictions about which aspects of brain health will be modulated by fitness levels in explaining older adults’ abilities to resolve tip-of-the-tongue states. It is not clear if one, multiple, or indeed any of the models align with age-related differences in tip-of-the-tongue states. Our objective in the current contribution, therefore, is twofold: 1) to provide a holistic account of tip-of-the-tongue states in ageing, incorporating brain health and cardiorespiratory fitness measures, and 2) to embed our findings within and provide support for existing frameworks of neurocognitive ageing. To test the suppositions of these models in a single contribution requires the capture of multimodal data, ideally in a single design. Here, for the first time, we provide such a contribution.

## Methods

Data and stimulus materials as relevant to the present report can be found here: https://osf.io/d7aw2/files/osfstorage.

### Participants

The sample comprised 73 (36 female, 37 male) community-dwelling, right-handed, healthy older adults aged 60-81 (M = 65.51, SD = 4.94). Neuroimaging data were collected in a single session; a cardiorespiratory fitness measure was performed on a different day (on average 9 days after the neuroimaging data). Participants underwent physical and cognitive screening which included a 12-lead resting electrocardiogram (ECG), blood pressure measures at rest, and the Montreal Cognitive Assessment (MoCA, Nasreddine et al., 2005). Participants were excluded if they presented with an abnormal ECG trace, were hypertensive, and/or scored ≤23 on the MoCA (Carson, Leach, & Murphy, 2018). We only included participants who reported to partaking in less than 150 minutes of exercise per week, did not suffer from a severe health condition (e.g., cancer, dementia, cardiorespiratory disease, etc.), and were non-smokers. All participants provided informed consent and were compensated £10.00 (GBP) per hour of participation. All were British-English monolinguals with no speech or language disorders. We did not collect race/ethnicity data. The research was conducted at the Centre for Human Brain Health and School of Sports, Exercise, and Rehabilitation Sciences, both at the University of Birmingham, UK. The study was granted institutional ethics approval (University of Birmingham, ERN 20_1107) and complied with the Declaration of Helsinki.

### Tip-of-the-tongue states as measured by a definition task

In the MRI scanner, participants completed a task whereby definitions were presented in text format alongside three response options: (1) Know, (2) Don’t Know, and (3) Tip-of-the-tongue (TOT). Respondents were instructed to select *Know* if they knew the word to which the definition was referring, *Don’t Know* if they did not know, and *TOT* if they had a tip-of-tongue experience. The proportion of tip-of-the tongue responses was calculated by dividing the number of true tip-of-the-tongue states (indication and verification of a tip-of-the-tongue state, see below) by the total number of trials observed. See Table 1 for example stimuli and Figure 1 for a task schematic.

**Figure 1.**
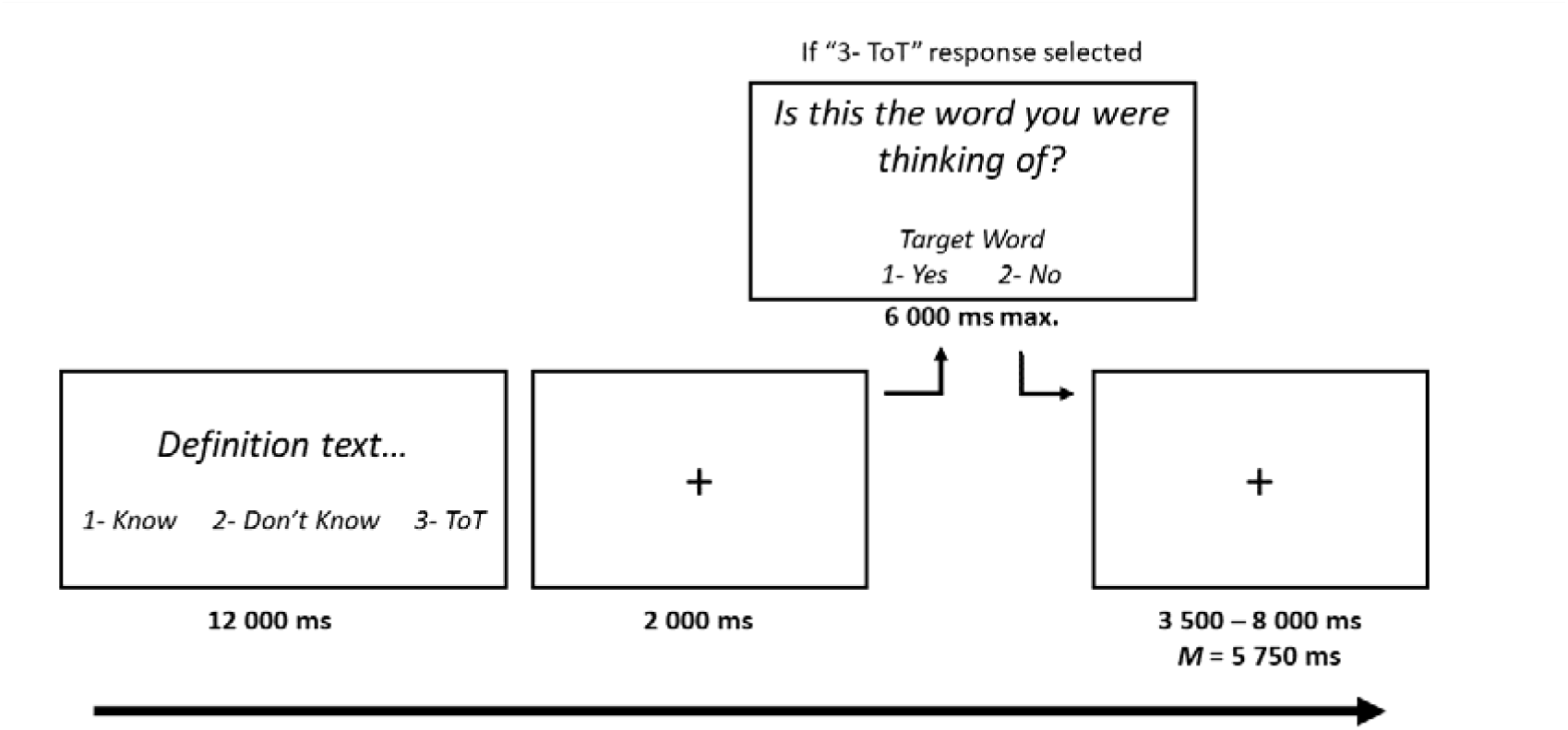
Schematic of the tip-of-the-tongue task. Trial lengths varied depending on the volume of tip-of-the-tongue responses (which triggered an additional verification slide that was not included for Know or Don’t Know responses). TOT = tip-of-the-tongue.

**Table 1.** Examples of definitions and their respective target words.

| Definition | Correct Answer |
| --- | --- |
| The termination of the existence of a particular kind of organism or species | Extinction |
| A word borrowed from German that means the "spirit of the times" | Zeitgeist |
| The boy king of Egypt whose tomb was discovered largely intact in 1922 | Tutankhamun |

A true tip-of-the-tongue was recorded only when participants indicated they were experiencing a tip-of-the-tongue state *and* pressed YES when asked “Is this the word you were thinking of?” in a subsequent verification slide (where participants pressed NO in response to the verification slide, these trials were considered in the same manner as Don’t Know responses). The definition texts were presented for a fixed duration of 12000 ms which was followed by a 2000 ms ISI and, in the case of tip-of-the-tongue responses, verification slides were presented for a maximum of 6000 ms (or sooner with button press). A jittered ITI was presented for an average of 5750 ms (range 3500 – 8000 ms; 500 ms increments). A total of 200 unique definitions were presented across four scanning blocks (50 per block); correct answers (targets) pertaining to definitions were arranged so that the number of proper and common nouns, syllable, phoneme, and letter count, as well as word frequency were counterbalanced across the four experimental blocks. The task was written in PsychoPy (v2021.1.4) and text was presented in white Open Sans font on a grey (normalised RGB 0.5, 0.5, 0.5) background projected onto a screen using a ProPixx visual projector system (1440 Hz refresh rate) and viewed by a mirror-system attached to the MRI headcoil. An MRI-compatible button box was used to register Know, Don’t Know, and TOT responses. Before entering the scanner, participants performed a practice run of the task with 12 definitions not included in the experimental stimuli—during this practice phase, an explanation of tip-of-the-tongue states was provided to ensure participants understood the task. There was no limit on practice; participants could repeat the practice task as many times as they wished. The set-up and parameters of the out-of-scanner practice task was identical to the in-scanner experimental task.

Behaviourally, from 200 stimuli presented across four fMRI runs, an average of 59.94% (SD = 13.52) of responses were recorded as Know, 23.19% as Don’t Know (SD = 10.98), and 16.87% (SD = 6.62) as TOT. The current proportion of tip-of-the-tongue states is comparable to what is typically reported in the ageing literature (e.g., Huijbers et al., 2017; Shafto et al., 2010).

### Cardiorespiratory Fitness

Participants completed an incremental exercise test on a treadmill (Pulsar 3p, H/P/Cosmos, Germany). Respiratory gases (V□O_2_; oxygen consumption, V□CO2; carbon dioxide production) were recorded continuously using a facemask (7450 V2, Hans Rudolph, USA) and metabolic cart (JAEGER Vyntus CPX, Vyaire, USA), as was heart rate and rhythm using a 12-lead ECG (Cardiosoft, Vyaire, USA). Participants warmed up on the treadmill before completing 4-min walking stages with a 1-min rest period between each stage. Treadmill speed started and remained at 3.8 km/h until either all elevation stages were completed (7 possible stages: 4, 7, 10, 13, 16, 19, and 20%) or individual lactate threshold was reached. Finger-prick blood lactate was measured during each rest period (Biosen C-Line, EKF Diagnostics, United Kingdom). If all elevation stages were completed, 4-min stages continued with speed increasing 0.5 km/h per stage until lactate threshold was reached or until volitional exhaustion. If lactate threshold was reached, participants immediately began 1-min stages where just the speed increased 0.5 km/h per stage until volitional exhaustion (rest periods were removed).

Participants were asked to exercise to volitional exhaustion unless halted by the researcher due to ECG abnormalities, injury, or the presence of a plateau in heart rate or V□O_2_. Cardiorespiratory fitness was measured as peak oxygen consumption (V□O_2_peak) recorded during the treadmill test, determined as the mean of the two highest 30 s V□O_2_ intervals recorded. For participants who only completed a sub-maximal test (N = 8), V□O_2_peak was predicted using sub-maximal V□O_2_ and heart rate data acquired from three of the first possible six stages using a linear regression where y = HRage-pred (age-predicted maximal HR, 220-age) (for more information, see Feron et al., 2024).

### Neuroimaging data acquisition and analyses

All neuroimaging data were acquired using a 3-T Siemens PRISMA system with a 32-channel head-coil at the Centre for Human Brain Health, University of Birmingham, UK. The scan sequence comprised (in order): T1-weighted anatomical, tip-of-the-tongue task-based fMRI, and pseudo-continuous arterial spin labelling (pcASL) scans. The total session lasted approximately 150 minutes (with ∼100 minutes in-scanner time).

*Brain structure*. A T1-weighted 3D-structural MRI (GRAPPA) was acquired with the following parameters: repetition time (TR) = 2000 ms, echo time (TE) = 2.01 ms, inversion time (TI) = 880 ms, flip angle = 8 deg, FOV = 256 x 256 x 208 mm, voxel dimension (resolution) = 1 mm isotropic, GRAPPA factor = 2; with a total acquisition time of 4 min and 54 s. Brain extraction was initially performed using FSL’s brain extraction tool (BET v2.1) but the standard Robust method performed poorly. Therefore, each participant’s anatomical image was manually extracted using the bias field and neck cleanup utility within FSL-BET (see Popescu et al., 2012); fractional intensity threshold, threshold gradient, and centre of gravity values were manually adjusted (where necessary) for optimal results. Brain-extracted images were then preprocessed and analysed with the standard FSL-VBM/FSL-ANAT pipelines. Sex, BMI, and intracranial volume (ICV) were included as covariates in the VBM analysis, where ICV was estimated using an atlas-based estimation procedure consistent with ENIGMA protocol (http://enigma.ini.usc.edu/). In terms of the VBM analysis, all images were first segmented into grey matter, white matter and CSF, while also correcting for spatial intensity variations (bias field or RF inhomogeneities), using FMRIB’s Automated Segmentation Tool (FAST). Grey matter images were then non-linearly registered to standard space using FMRIB’s Nonlinear Image Registration Tool (FNIRT) and modulation was applied by multiplying each registered grey matter image by the Jacobian determinants of the warp fields derived from the non-linear registration step. At this stage, the images were then used as predictors in the commonality analysis (i.e., these images constituted the ‘Structure’ variable). Separately, the remainder of the full standard FSL-VBM pipeline was then ran with a design matrix that included age, sex, ICV, and BMI.

*Cerebral Perfusion*. To measure CBF, a pcASL sequence with 3D GRASE readout (Wang et al., 2013) was used, see also Acknowledgements. ASL imaging parameters were: echo time (TE) = 30.56 ms, repetition time (TR) = 4100 ms, in-plane resolution = 3.5 mm^2^, slice thickness = 3.5 mm, transversal slices = 32, field of view (FOV) = 224 mm, labelling duration = 1508 ms, no background suppression or vascular crushing, and post-label delays (PLDs) = 200, 975, 1425, 1850, 2025, 2150, 2250, and 2300 ms. Four volumes of data were acquired for PLDs of 200 – 2250 ms and 12 volumes for 2300 ms. PLD times and number of volumes acquired were optimised according to Woods, Chappell, and Okell (2019). Slices were positioned axially from the motor cortex and angled anterior-posterior in line with the participant’s anterior-posterior commissure (ACPC). A base equilibrium, M_0_, scan was acquired using these same parameters with the PLD set to 2000 ms and no inversion pulses. Acquisition time was ∼17 minutes. The previously described T1-weighted anatomical images were used for differentiation and segmentation of grey and white matter so that resting CBF in only grey matter could be assessed.

ASL data were processed using the Oxford ASL toolbox (https://oxasl.readthedocs.io/en/latest/) which uses the FSL FABBER ASL package and Bayesian Inference to invert the kinetic model for ASL MRI (BASIL) to compute CBF and ATT (i.e., arterial transit time) maps (Chappell et al., 2008; Groves et al., 2009; Woolrich et al., 2006). Parameters input to the kinetic models to estimate CBF and ATT were: bolus duration = 1.508 s, tissue T1 = 1.3 s, arterial blood T1 = 1.65 s, labelling efficiency = 0.85, inversion times = 1708, 24838, 29338, 33588, 35338, 36588, 37588, and 38088 ms; all other input parameters were kept with default settings appropriate to pcASL acquisition. Partial volume error correction and adaptive spatial smoothing of the perfusion maps was performed using default settings in oxford_asl (Chappell et al., 2011; Groves et al., 2009).

Participants with abnormal post-label delay maps, often caused by motion but also due to incidental findings such that perfusion in part of the brain could not be detected, were excluded (N = 6). Grey matter masks were thresholded to ensure only voxels containing primarily grey matter were included. Areas within masks containing incorrect assignment to grey matter, primarily around the eyes and nasal cavity, were manually removed (N = 7). ATT measures are less common in MRI perfusion measures, as often only one PLD is used. In our work, per-participant ATT measures were used to correct CBF for ATT-differences, improving CBF estimation accuracy. ATT itself was used an outcome measure in another paper (Feron et al., 2024).

*Brain function.* Four fMRI blocks (50 definitions per block) were run with an echo-planar imaging sequence with the following parameters: TR = 1500 ms, TE = 35 ms, slice thickness = 2.5 mm (2.5 mm isotropic voxel size), FOV = 210 mm, flip angle = 71 deg, phase encoding direction = A >> P, and interleaved slices. The total number of volumes per run varied according to the number of tip-of-the-tongue states reported; typically, ∼610 volumes were collected per run (∼16 min/run).

Both individual- and group-level data were processed using FEAT (fMRI Expert Analysis Tool) in FMRIB’s Software Library (FSL, www.fmrib.ox.ac.uk/fsl) v6.0.1. Registration to the high-resolution T1-weighted structural (BBR [Boundary-Based Registration, Greve & Fischl, 2009]) and standard space images (MNI152 T1 2mm; 12 DOF transformation) was performed using FLIRT. Registration from high-res structural to standard space was then further refined using FNIRT nonlinear registration. After registration, the following pre-statistics processing was applied: motion correction using MCFLIRT (Jenkinson et al., 2002), slice-timing correction (interleaved) using Fourier-space time-series phase-shifting, spatial smoothing using a Gaussian kernel of FWHM 5 mm, grand-mean intensity normalisation of the entire 4D dataset by a single multiplicative factor, highpass temporal filtering (Gaussian-weighted least squares straight line fitting, with sigma = 50.00 s). The time-series were analysed using FMRIB’s Improved Linear Model tool with local autocorrelation correction (Woolrich et al., 2001) and Z (Gaussianised T/F) statistics images were thresholded non-parametrically using clusters determined by Z>3.1 and a corrected cluster significance threshold of p = .05. Whole-brain functional univariate analyses were computed within the general linear model using a multi-level mixed-effects design. Each fMRI run was modelled separately at the single-participant level. Trial-by-trial modelling commenced at trial onset and concluded at participant button press within only the definition presentation phase of the task. Participant responses, and therefore statistical modelling, were not captured once definitions disappeared from screen (BOLD signal during ISI/TOT verification slide was not modelled). Trials where participants failed to make a response were discarded. Each condition of interest or exploratory variable (TOT, Know, Don’t Know) was convolved using a double-gamma hemodynamic response function. A second-level analysis combined first-level contrast estimates per participant using a fixed-effects model, and higher-level activation (third-level analysis) was computed using a mixed-effects model using FSL’s FLAME1 (FMRIB’s Local Analysis of Mixed Effects) tool (Woolrich et al., 2009).

As the number of events (tip-of-the-tongues) varied from participant to participant, there was an imbalance in our statistical design. For example, it could not be assumed that participants producing <5% tip-of-the-tongue rates were statistically comparable—in the BOLD analysis—to those scoring >20% because of the mismatch in the number of timepoints/events. To address this, we additionally ran our higher-level analysis with FLAME1+2 modelling, which is a Bayesian-based method that estimates the inter-participant random-effects component of the mixed-effects variance by using Markov Chain Monte Carlo (MCMC) randomisation to produce a more precise estimate of the true random-effects variance and degrees of freedom on a voxel-by-voxel basis (see Jenkinson et al., 2012; Smith et al., 2004; Woolrich et al., 2009). Overall, FLAME methods are robust to unbalanced designs and use MCMC to better estimate the distribution of effects when sample sizes are low or variances are unequal, providing a more conservative and accurate inference framework than simpler models like ordinary OLS or fixed-effects-only approaches. We found that the results of the FLAME1+2 approach did not meaningfully alter what was yielded from the original FLAME1 approach, and therefore we retained the output of FLAME1 and fed this into a commonality analysis.

### Commonality Analysis

To explore the unique and shared contribution of tip-of-the-tongue-related functional activity, grey matter density, cerebral blood flow (CBF), cardiorespiratory fitness (V□O_2_peak) and age on behavioural tip-of-the-tongue rates, we conducted a commonality analysis (Kraha et al., 2012; Nimon et al., 2008). Commonality analysis partitions the variance accounted for by all predictors in multiple linear regression into variance unique to each predictor and common variance shared between each possible combination of predictors. As such, unique effects specify the (orthogonal) variance accounted for by one predictor above and beyond that accounted for by other factors in the model, while common effects show the combined variance shared between correlated predictors. Coefficients of common effects can show how much variance is accounted for in the outcome variable mutually by two or more correlated predictors. More precisely, in terms of interpretation, a positive commonality coefficient between two predictors does not necessarily mean that both predictors individually show positive relationships with the dependent variable. Rather, it reflects that the shared variance between those predictors *adds positively* to model’s total explained variance. It means that, even though the predictors may relate to the outcome in different directions, the pattern of how they relate to each other, and the outcome variable supports the prediction. This is similar to principal component analysis, where two variables can have loading values with different directions. Conversely, a negative shared coefficient can occur in the presence of suppression or when some of the correlations among predictor variables have opposing signs (Kraha et al., 2012; Nimon et al., 2008; Pedhazur, 1997). For a more comprehensive account of subjecting neuroimaging and behavioural data to commonality analysis, please refer to Wu et al. (2023). The commonality toolbox used in the present analysis is available at https://github.com/kamentsvetanov/CommonalityAnalysis. Significant clusters related to effects of interest (as outlined above) were reported, as identified with nonparametric testing using 1000 permutations and threshold-free cluster enhancement (TFCE) with significance level of a = .05 (Smith & Nichols, 2009).

Sex differences were not of primary interest in the current contribution but are controlled for because they may covary with multiple components of our data set, including total brain volume (which affects grey matter quantification) and cardiorespiratory fitness levels (males typically show higher V□O_2_ scores). Sex was included in the modelling as a binary covariate of no interest. Head motion has been shown to lead to spurious outcomes in functional neuroimaging data (e.g., Power et al., 2012) and has been linked with out-of-scanner demographic and lifestyle factors such as age (Madan, 2018) and BMI (Beyer et al., 2021), which in turn correlate with fitness. To control for this, head motion was included as a covariate of no interest in the commonality modelling. Head motion data were derived from FSL MCFLIRT reports where motion was indexed as relative displacement (in mm), which is a measure of the mean head movement in relation to the subsequent volume. These scan-specific values were calculated per FMRI run, and then averaged across runs to compute overall head motion such that a single value represented each participant’s total head motion. Recall, however, that the initial pre-commonality fMRI analysis was already motion corrected as part of the standard pre-processing of functional MRI data.

In addition to the primary model (see Figure 3), a further model was run where the following variables were included as covariates: education and vocabulary size (depicted in Figure 3 is the primary model with initial covariates, as well as the additional model with two added covariates). Education data were captured as part of the initial screening procedure and vocabulary was captured via an online task where participants were required to select the correct synonym or antonym of a target word. Education was categorised according to the British (English) education system: “Compulsory” = Schooling completed up to and including the age of 16 (N = 21); “College/FE” = Post-16 education including A-Levels and Further Education (N = 25); “Undergraduate” = Undergraduate qualification at university level (N = 14); “Postgraduate” = Postgraduate qualification at university level (N = 13). As education has widespread influences in linguistic and non-linguistic cognitive function (Zahodne, Stern, & Manly, 2015), is a purported component of cognitive reserve (Tucker & Stern, 2011), and is associated with cardiovascular health in ageing (Schultz et al., 2018), education was included in the modelling.

## Results

The data in the current report were acquired as part of a larger intervention study titled Fitness, Ageing, and Bilingualism (www.fab-study.com), which was detailed in a single preregistration on the OSF: https://osf.io/d7aw2. The preregistration contains a multitude of hypotheses and analyses plans that form the basis of other, separate reports (e.g., Fernandes et al., 2024; Fosstveit et al., 2024; Markiewicz et al., 2024). No one specific hypothesis in the preregistration is related to the current contribution, though several hypotheses related to Research Question 3 pertain to the relationship between fitness-levels and each of the brain-based measures included. All datatypes and several analysis steps used in the present report were specified in this preregistration.

An overview of sample characteristics is provided in Table 2.

**Table 2.** Overview of core participant sample characteristics.

| Variable |  | *M* | *SD* | Range |
| --- | --- | --- | --- | --- |
| Age |  | 65.51 | 4.94 | 60—81 |
| V̇O<sub>2</sub>peak |  |  |  |  |
|  | Male | 30.23 | 3.83 | 23.30—36.85 |
|  | Female | 24.80 | 3.43 | 17.00—31.35 |
|  | All | 27.55 | 4.53 | 17.00—36.85 |
| BMI |  |  |  |  |
|  | Male | 27.88 | 3.47 | 21.50—35.60 |
|  | Female | 26.30 | 3.62 | 20.20—34.10 |
|  | All | 27.10 | 3.61 | 20.20—35.60 |
|  |  | *N* | % |  |
| Gender |  |  |  |  |
|  | Male | 37 | 51 |  |
|  | Female | 36 | 49 |  |
| Education |  |  |  |  |
|  | Compulsory education | 21 | 29 |  |
|  | Further education | 25 | 34 |  |
|  | Undergraduate degree | 14 | 19 |  |
|  | Postgraduate degree | 13 | 18 |  |

Note: V̇O<sub>2</sub>peak (mL/kg/min). BMI = Body Mass Index (kg/m<sup>2</sup>). Education: Compulsory = mandatory education under the British (English) system; Further = Post-16 study after secondary education (and prior to university); Undergraduate = bachelor's degree or equivalent; Postgraduate = qualification undertaken after a bachelor's or undergraduate degree.

Figure 2 depicts the key bivariate relationships in our healthy older adult sample, corroborating findings from prior studies. With increasing age, the incidence of tip-of-the-tongue states increased (Panel A) and cardiorespiratory fitness levels (measured as V□O_2_peak) decreased (Panel B). Age did not significantly impact on overall cerebral blood flow (Panel C) (see Feron et al., 2024 for more detailed discussion). FMRI-BOLD analysis of the Tip-of-the-tongue > Know contrast produced significant clusters of activation within the precuneus and angular, cingulate, and frontal gyri (Panel D, and Table 3, Huijbers et al., 2017). Voxel-based morphometry analysis of whole-brain grey matter density showed significant and widespread age-related atrophy consistent with non-pathological ageing (Panel E, e.g., see Giorgio et al., 2010).

**Figure 2.**
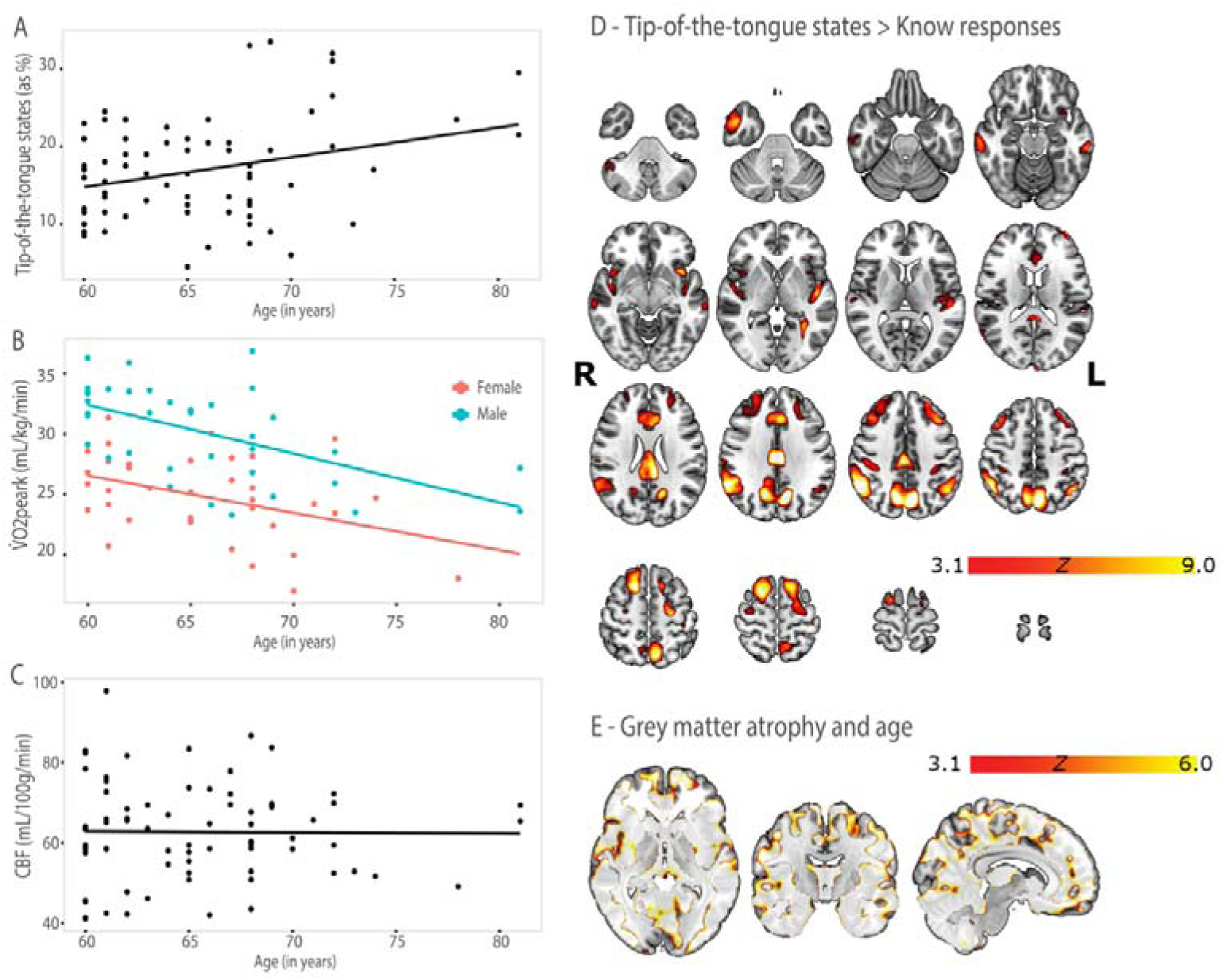
Bivariate analyses of the precursors to commonality analysis. A. Tip-of-the-tongue by age relationship (*r* = .288, *p* = .014). B. O_2_peak by age as a function of sex (Males *r* = −.566, *p* < .001, Females *r* = −.407, *p = .*014). C. Cerebral blood flow by age (*r* = −.011, *p* = .926). D. Significant clusters (determined by Z>3.1 and a corrected cluster significance threshold of p = .05) from the Tip-of-the-tongue > Know contrast. Slices, presented in MNI space, begin (top left) at z = −40 and conclude at z = 80 (bottom right), with each progressive slice representing an increment of +8. E. Volumetric analysis depicting grey matter atrophy by age (voxel-based morphometry analysis with 10,000 permutations using threshold-free cluster enhancement). All brain images are presented in radiological convention.

**Table 3.**
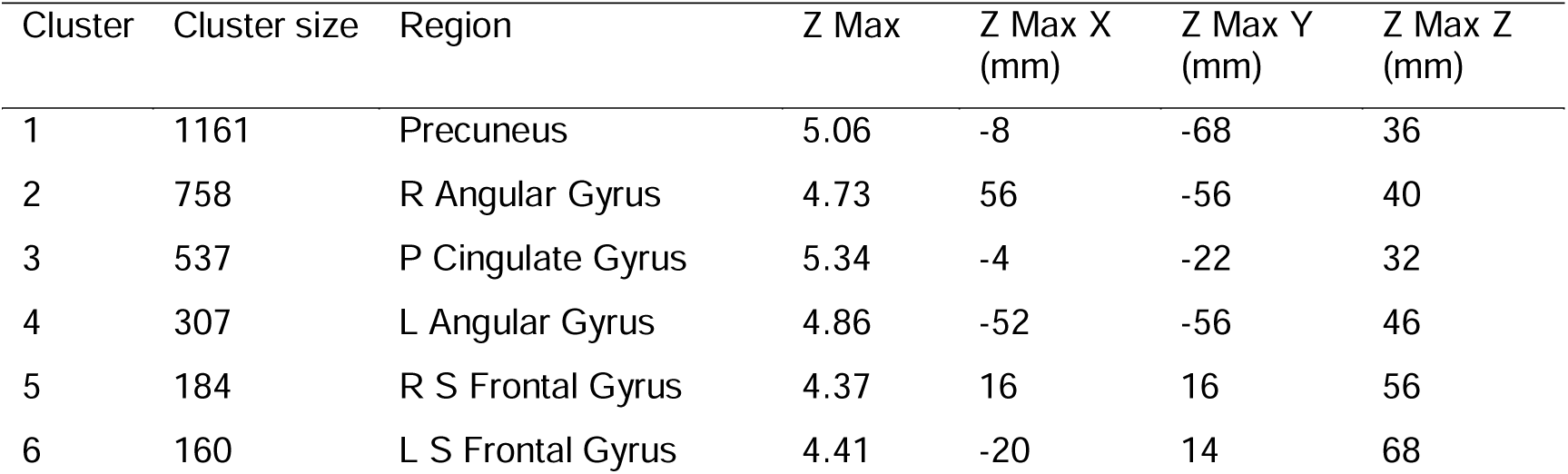
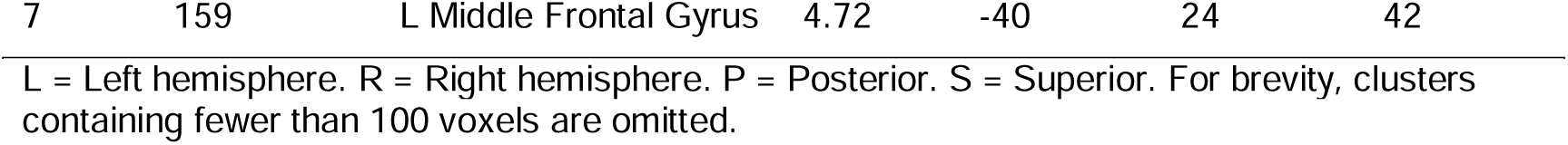
Coordinates (MNI), size, and z value of brain regions showing significant tip-of-the-tongue activation (Tip-of-the-tongue > Know).

Next, we analysed all data modalities with one joint approach using commonality analysis, to determine a parsimonious combination of factors—which were all individually theoretically-motivated, as shown above—in explaining healthy older adults’ tip-of-the-tongue rates. Figure 3 summarises the main components of the present multimodal dataset, feeding into a commonality analysis. The term *functional* refers to functional MRI data derived from the contrast in BOLD signal between TOT and Know responses, *structure* or *grey matter* are used interchangeably to refer to the quantification of whole-brain grey matter volume, and *fitness*, *cardiorespiratory fitness*, and V□*O_2_peak* are used to refer to cardiorespiratory fitness. Our core model of interest included the three brain-based datatypes, a measure of cardiorespiratory fitness, and age. As sex and in-scanner head motion can affect brain measures (Ruigrok et al., 2014) and/or are related to core predictors, these variables were additionally entered into the model as covariates of no interest.

**Figure 3.**
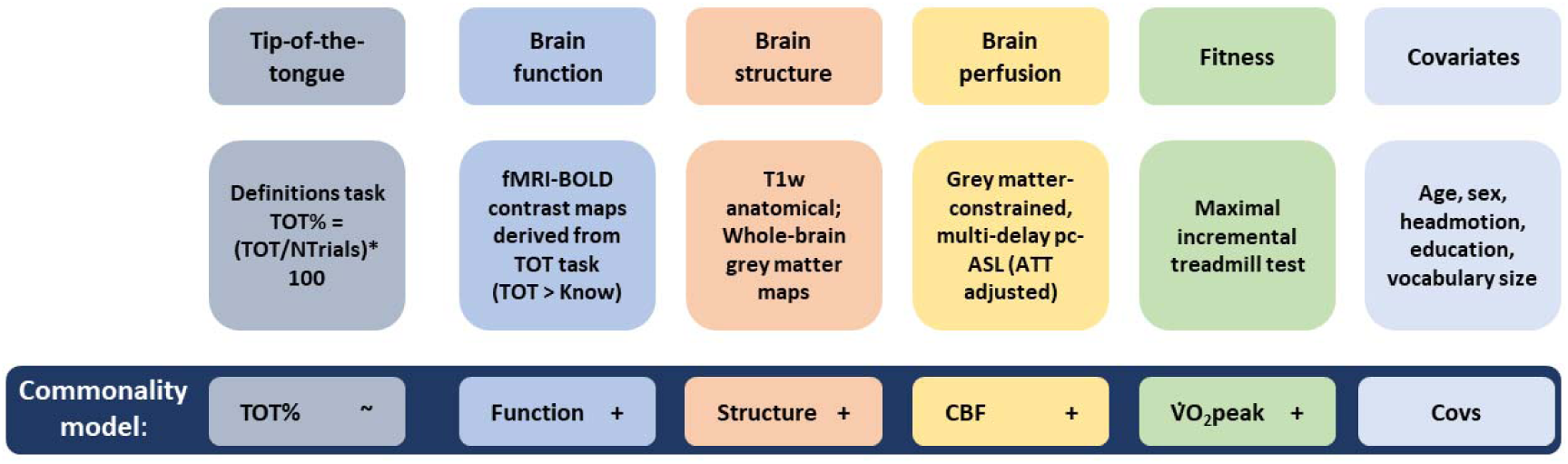
Summary of the main components of the commonality analysis used to analyse the current multimodal data. For information on how each component was measured and modelled, please refer to Methods. TOT = tip-of-the-tongue. CBF = cerebral blood flow. pc-ASL = pseudo-continuous arterial spin labelling. ATT = arterial transit time.

Complementing the bivariate functional analyses, commonality was found between age and function in explaining variance of tip-of-the-tongue rates in regions including the cerebellum, middle temporal gyrus, inferior frontal gyrus (pars triangulars), and superior frontal gyrus (medial part). See Table 4 and Figure 4B for details of significant contrasts and their associated coefficients. Note that the regions reported for the age and function common effect show some degree of dissimilarity relative to the regions reported previously in the pre-commonality analysis. The reasons for this are twofold. First, only regions that expressed commonality in age *and* function and survived stringent thresholding are reported for the commonality analyses; and second, clusters containing fewer than 150 voxels were not reported in the pre-commonality analyses for the sake of brevity but are reported here due to overall smaller effects.

**Figure 4.**
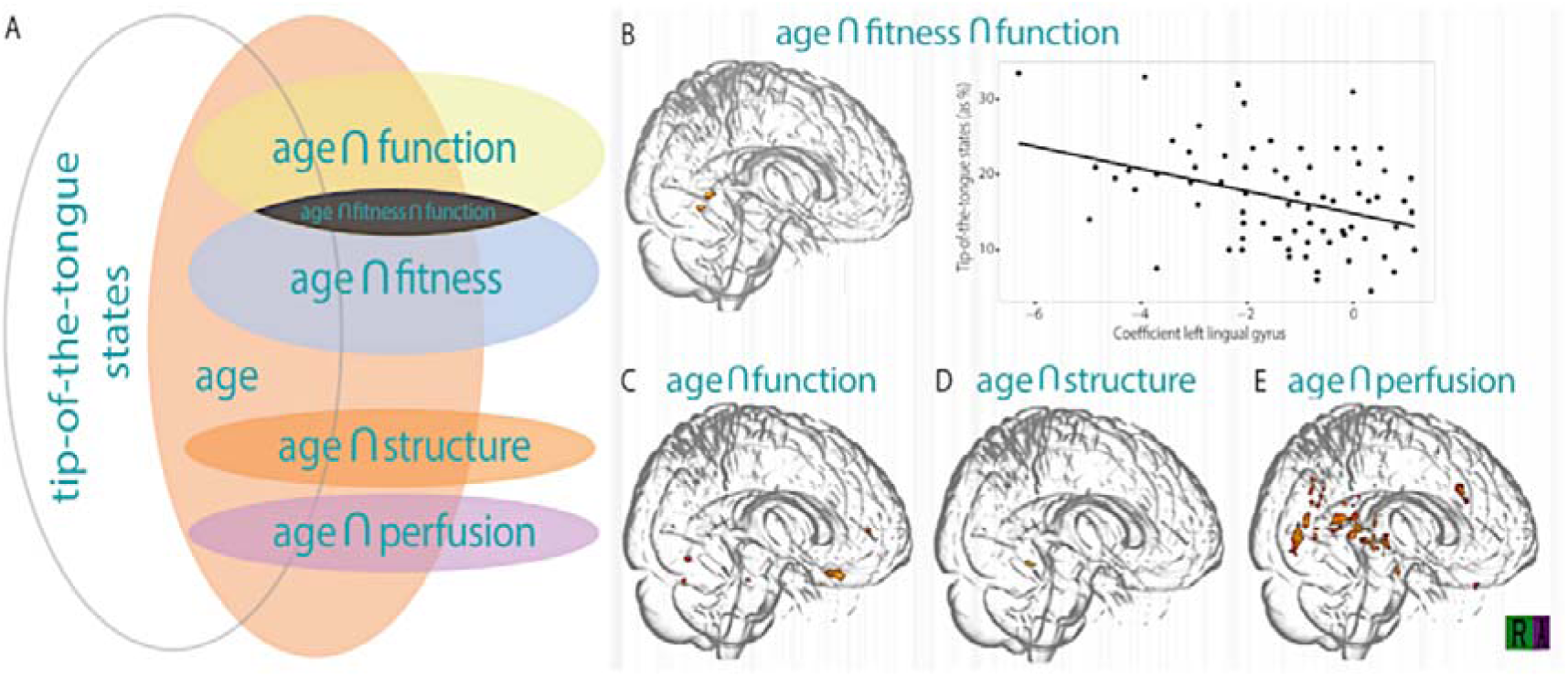
A: Euler diagram depicting the relationships, as established by the commonality analysis, where, for example, the intersection tip-of-the-tongue states on the one hand with age and function on the other hand, depicts that the commonality between age and function explains a significant portion of the variance in tip-of-the-tongue states. The size of the ellipses is only illustrative, it is not indicative of effect size. Our key finding is the common effect of age, fitness and function (i.e., black intersection) accounted for a significant portion of variance in tip-of-the-tongue states (which is depicted further in Panel B). Age-associated structural atrophy and perfusion in regions other than those showing functional differences also accounted for significant portions of variance in tip-of-the-tongue states (i.e., orange and purple intersections respectively) but these did not share commonality with fitness. B. Three-dimensional render depicting the age, function, and fitness common effect including a scatter presenting the relationship between this contrast and its intensity in the region of interest analysis (lingual gyrus). C. Age by function common effect. D: Age by structure common effect. Age and structure common effect did not converge with function or fitness. A significant age by fitness common effect was observed but is not visualised here. Heatmaps are indicative of *t* values, with brighter colours representing stronger effects (*t* range 1.80 – 3.00). Commonality effects were observed using threshold-free cluster enhancement (TFCE) correction. ∩ = intersection (commonality between stated predictors).

**Table 4.** Coordinates (MNI), size, *t,* and *p* values of significant commonality effects and their respective brain regions.

| Contrast | Cluster size (k) | Cluster | $t$ | $p$ | x | y | z |
| --- | --- | --- | --- | --- | --- | --- | --- |
| Age.Function | 67 | R. Cerebellum Crus VI | 1.897 | .009 | 32 | -56 | -24 |
|  | 55 | L. Middle Temporal Gyrus | 1.921 | .009 | -56 | -4 | -20 |
|  | 40 | L. Inferior Frontal Gyrus - pars triangulars | 1.848 | .010 | -50 | 16 | 0 |
|  | 45 | R. Superior Frontal Gyrus - Medial Part | 1.944 | .009 | 10 | 64 | 4 |
| Age.Function. O<sub>2</sub>Peak | 27 | L. Lingual Gyrus | -1.842 | .002 | -6 | -64 | 0 |
| Age.Structure | 9 | R. Middle Temporal Pole | 2.196 | .001 | 46 | 12 | -38 |
| Age.CBF | 46 | L. Cerebellum Crus I | 1.810 | .025 | -24 | -70 | -28 |
|  | 12 | L. Cerebellum Crus IX | 2.077 | .015 | -18 | -42 | -48 |
|  | 10 | R. Cerebellum Crus IX | 2.047 | .014 | 8 | -50 | -44 |

A unique effect of age was also present but not reported in the above table because an isolated age effect is uninformative and was not hypothesised. Note: R = Right. L = Left. CBF = Cerebral blood flow.

Interestingly, and key to the present research question, we also found commonality between age, function, and VO _2_peak in explaining age-related tip-of-the-tongue variability. More precisely, the negative shared coefficient suggests a suppressor effect. While age is positively associated with tip-of-the-tongue states, this effect is attenuated in individuals with higher fitness and greater BOLD activation. The negative coefficient reflects that the combination of high VOLpeak and strong functional recruitment in some older adults offsets part of the age-related variance in tip-of-the-tongues. While these effects were not widespread, with the effect being localised to the left lingual gyrus, the finding was robust across multiple iterations of commonality modelling (see below for more information on models including education and vocabulary size). It is worth noting, however, function did not produce a shared effect with brain structure or perfusion, suggesting that the functional effects are over and above any differences in atrophy and/or perfusion.

A positive shared effect was also found between age and structure in the middle temporal pole. This indicates that age-related structural decline helps explain increased tip-of-the-tongues in older adults. Although grey matter volume in this region was not uniquely associated with tip-of-the-tongues (i.e., there was no unique effect of Structure), its shared variance with age contributes to explaining age-related difficulty in word finding.

A positive coefficient between age and perfusion indicates that age-related changes in CBF contribute to increased tip-of-the-tongues. While age and CBF were not significantly correlated at the bivariate level, their shared variance nevertheless explains part of the relationship between age and word finding failures. This again suggests that indirect neural mechanisms related to vascular ageing play a role in the behavioural manifestation of tip-of-the-tongues.

The distinct effects of brain structure, perfusion, and function suggest that the mechanisms by which these three indices of brain health contribute to tip-of-the-tongue in healthy ageing are independent.

In addition to the core model of interest outlined above, we ran additional analyses which included the variables education and vocabulary size. We considered these variables as indicators of crystallised intelligence and found a moderate-to-strong positive correlation between them (*r* = .638, *p* <.001). However, neither education nor vocabulary were significant—uniquely or jointly with other predictors—in explaining variance in tip-of-the-tongue occurrences.

## Discussion

Here, for the first time, we bring together multiple brain-health measures and a gold-standard assessment of fitness to explain the differences of word-finding difficulties and more broadly cognitive decline in older adults. Commonality analyses revealed that while age is positively associated with tip-of-the-tongue states, this effect is attenuated in individuals with higher fitness and greater BOLD activation within the functional network related to tip-of-the-tongues. Further, age-associated changes in brain structure and age-related cerebrovascular differences in brain regions other than those showing functional differences accounted for the variance in tip-of-the-tongue state incidences.

Our findings can be interpreted in the light of key neurocognitive models of ageing, such as maintenance, compensation, or reserve. We found no evidence supporting maintenance and reserve indicators: neither grey matter nor perfusion, nor indicators of reserve such as education, showed shared effects with task activity on predicting tip-of-the-tongue behaviour. We did, however, find evidence that could be interpreted as compensation: there was a shared effect of functional recruitment, cardiorespiratory fitness, and age predicting tip-of-the-tongue incidence. The unison of these factors in explaining variance in age-associated tip-of-the-tongue states suggests that both fitness and age modify the functional recruitment of word-finding areas in the brain. While some previous studies found evidence of fitness levels contributing to altered brain function (Douw et al., 2014; Prakash et al., 2011; Voss et al., 2016; Wong et al., 2015), these studies did not formally explore the connections to cognitive performance as examined in the present study. Future studies could use our present holistic approach integrating multimodal brain-health combined with lifestyle measures to examine other indicators of language abilities as well as other domains of cognition. Our findings indicate that older adults with higher fitness levels display distinct behaviourally-relevant brain activity compared to their less fit counterparts, providing a brain-based explanation for the bivariate relationship observed in previous research between fitness and tip-of-the-tongue states (Segaert et al., 2018).

Our findings also build upon previous work which has found that atrophy (Shafto et al., 2007) was associated with age-related tip-of-the-tongue rates. Specifically, age-related increases in tip-of-the-tongue states have been associated with atrophy in the left insula, a region implicated in phonological production (Shafto et al., 2010). Similarly, in the current contribution, we report commonality between age and brain structure, albeit not in the insula region but in the right middle temporal pole. Further to functional and structural brain effects, we also observed that individual differences in cerebral perfusion, as a function of age, explained variation in tip-of-the-tongue states. The literature on the association between age-related differences in cerebral blood flow, cognition, and wider brain health is unresolved (for a review on perfusion and cognition in ageing, see Ogoh, 2017): while some accounts report perfusion maintenance in older adulthood supporting brain function and cognition (De Vis et al., 2018; Wu et al., 2023), others have shown mixed results (Espeland et al., 2018; Leeuwis et al., 2018; Poels et al., 2008). With our data, we bring further clarity by extending the literature to show that cerebral blood flow is an important contributor to age-related tip-of-the-tongue states.

We focused on tip-of-the-tongue variability as an example of cognitive decline in older age. Failure to produce known words is a common human experience; it often leads to frustration and disrupts the natural flow of verbal communication. While it is not unique to ageing, healthy older adults tend to display more tip-of-the-tongue occurrences relative to their younger counterparts. This age-related pattern cannot be explained by memory deficits alone—indeed, semantic and phonological features of the intended yet elusive word can often still be articulated (Brown, 1991; Brown & McNeill, 1966; Miozzo & Caramazza, 1997). Moreover, previous work has shown the independence of memory and word retrieval/production systems (Salthouse & Mandell, 2013). We demonstrate, for the first time, that tip-of-the-tongue variability in ageing can be potentially explained by variability in brain function, structure, and perfusion in addition to fitness, in one analytical framework, and provide a theoretical and methodological step forward in bringing together seemingly disparate streams of data to profile a recognisable, everyday age-related irritation. Our approach could be applied to other aspects of cognitive function which are relevant to characterising age-related cognitive decline.

Altogether, our commonality analysis provides nuanced insight into the mechanisms underpinning age-related increases in tip-of-the-tongue experiences. These findings highlight a key strength of commonality analysis: it can detect shared explanatory pathways that may not emerge through conventional univariate or multivariate regression approaches. In cognitive ageing research, where multiple predictors are often intercorrelated and related to behavioural outcomes in complex ways, this approach helps illuminate indirect and interactive mechanisms that contribute to cognitive change.

### Future directions

There are several potential extensions to the present work. Our cross-sectional study cannot directly show individual progression over time (i.e., the ageing process) and cannot rule out pre-existing lifelong, trait-level differences in functional recruitment, possibly limiting our ability to distinguish between neurocognitive ageing models. In future work, longitudinal studies are needed to clarify the conditions and sequences of events behind our findings. This could include longitudinal intervention work, as changes in lifestyle, for example, increasing fitness through changes in regular physical activity accumulated over time, would help to elucidate the framework we have set out in the present work. Moreover, while we examined brain activations and co-activations, we did not quantify functional connectivity (Bethlehem et al., 2020; Geerligs et al., 2017; Liu et al., 2023; Tsvetanov et al., 2016). Both aspects may change independently during cognitive ageing, requiring further investigation (Tsvetanov et al., 2018). We believe that our current contribution provides a sound theoretical basis and analytical framework to motivate future longitudinal designs.

Neurocognitive models of ageing are continually evolving, as are the ways in which these constructs are best studied (Stern et al., 2020). Future work could validate our findings using diverse approaches to define and assess these constructs. Our method of recording education data — a proxy for cognitive reserve—relied on self-reported categorical data with low variability. The lack of effect with education does not suggest that education is an unimportant variable in characterising cognitive reserve. Future studies could use more sensitive methods to measure education in combination with crystallised intelligence. Moreover, our understanding of what constitutes cognitive reserve is evolving. Factors such as physical activity (and resultant changes in fitness levels) or bilingualism may also contribute to individual variations in cognitive reserve (Cabeza et al., 2018).

The use of a cross-sectional design implies that mediation analyses techniques such as structural equation modelling (SEM) would not be appropriate to apply to the data. When applied to point-in-time, age-heterogenous data, SEM may not accurately capture relationships or capture individual differences over time, as its assumptions are often violated with such data (e.g., see Hofer et al., 2006; Imai, Keele, & Tingley, 2010; and Maxwell & Cole, 2007). Commonality analysis was the more suitable approach for this dataset. We also note that while our study included participants aged 60 to 81, 84% of participants were aged 60-69, 13% were aged 70-79, and only 3% were aged 80 or above. While this inequality in age distribution is understandable given the stringent nature of our inclusion criteria, it nonetheless means that the current analysis perhaps lacked sensitivity at the highest bands of age in our sample, and we were unable to test whether the reported effects differed across older age groups. We do, however, contend that chronological age is only one component that determines cognitive ageing; here, a much wider perspective on how the effect of ageing is potentially modified by cardiorespiratory fitness, and how this then downstreams into cognition, is provided. Nevertheless, sampling more widely in future work is advisable. Further, in the present work, individual participants’ arterial transit time (i.e., the time it takes blood to travel from an arterial input to a brain region) measures were used to the correct our cerebral blood flow assessment for arterial transit time-differences, improving the estimation accuracy of cerebral blood flow. The measure of arterial transit time could be added in future work to supplement the standard cerebral blood flow measure (see Feron et al., 2024). The inclusion of arterial transit time may highlight potential commonality with functional BOLD signal (e.g., if blood takes longer to reach an ‘activated’ area due to a prolonged arterial transit time, it may delay the onset of, and therefore distort, the BOLD signal response). Finally, both performance and BOLD contrast were derived from the same trial types, introducing potential statistical dependence. However, the effects were explained by the relationship with an independent third variable (fitness, i.e., shared effects between function and fitness), which helps contextualise the brain–behaviour link. Future work could consider trial-splitting with commonality analysis to better isolate unique variance components.

## Conclusion

In sum, our multi-modal approach, which integrates brain structure, function, perfusion, and fitness measures offers a useful framework for advancing research on cognitive ageing. We demonstrate that cardiorespiratory fitness is associated with brain function and contributes to cognitive performance in healthy older adults. The combination of higher cardiorespiratory fitness and functional recruitment in some older adults offsets part of the age-related variance in tip-of-the-tongues rates. Age-associated atrophy and perfusion in regions other than those showing functional differences furthermore accounted for variance in tip-of-the-tongues rates. These findings emphasize the importance of investigating associations between lifestyle factors such as respiratory fitness and brain function mechanisms, as they are potentially promoting compensatory mechanisms.

## Acknowledgements

This work was funded by the Norwegian Research Council (FRIPRO 300030). We thank Sue Sandys for help with creating the tip-of-the-tongue stimulus lists. We also thank Caroline Ratcliffe, Charnjit Sidhu, Dagmar Fraser, and Jan Zandhuis for structural and technical support. Bethany Skinner, Consuelo Vidal Gran, Nicolas Hayston, Rupali Limachya, Amelie Grandjean, Aoife Marley, Shi Miao, and Samuel Thomas helped with data collection. Finally, we thank the wonderful and committed participants that contributed to the FAB project. KAT was supported by the Guarantors of Brain (G101149) and the Alzheimer’s Society (Grant Number 602). We thank Danny Wang and the University of Southern California’s Steven Neuroimaging and Informatics Institute for the provision of the pcASL sequence used in this work which was provided through a C2P agreement with The Regents of the University of California.

The authors declare no competing interests.

